# A nanopore based chromosome-level assembly representing Atlantic cod from the Celtic Sea

**DOI:** 10.1101/852145

**Authors:** Tina Graceline Kirubakaran, Øivind Andersen, Michel Moser, Mariann Arnyasi, Philip McGinnity, Sigbjørn Lien, Matthew Kent

## Abstract

Currently available genome assemblies for Atlantic cod (*Gadus morhua*) have been constructed using DNA from fish belonging to the Northeast Arctic Cod (NEAC) population; a migratory population feeding in the cold Barents Sea. These assemblies have been crucial for the development of genetic markers which have been used to study population differentiation and adaptive evolution in Atlantic cod, pinpointing four discrete islands of genomic divergence located on linkage groups 1, 2, 7 and 12. In this paper, we present a high-quality reference genome from a male Atlantic cod representing a southern population inhabiting the Celtic sea. Structurally, the genome assembly (gadMor_Celtic) was produced from long-read nanopore data and has a combined contig size of 686 Mb with a N50 of 10 Mb. Integrating contigs with genetic linkage mapping information enabled us to construct 23 chromosome sequences which mapped with high confidence to the latest NEAC population assembly (gadMor3) and allowed us to characterize in detail large chromosomal inversions on linkage groups 1, 2, 7 and 12. In most cases, inversion breakpoints could be located within single nanopore contigs. Our results suggest the presence of inversions in Celtic cod on linkage groups 6, 11 and 21, although these remain to be confirmed. Further, we identified a specific repetitive element that is relatively enriched at predicted centromeric regions. Our gadMor_Celtic assembly provides a resource representing a ‘southern’ cod population which is complementary to the existing ‘northern’ population based genome assemblies and represents the first step towards developing pan-genomic resources for Atlantic cod.

## Introduction

Atlantic cod (*Gadus morhua*) is a commercially exploited high-fecundity fish with a wide geographical distribution extending over the North Atlantic Ocean from the nearly freezing waters in the Arctic to variable high temperatures typical of the southern extremities of the species’ Eastern Atlantic distribution (Mieszkowska et al. 2009; Righton et al. 2010; Morris et al. 2018). It has been proposed that increases in water temperatures associated with global warming will see Atlantic cod spread northwards and occupy larger areas of Barents Sea, while southern populations will decline and possibly disappear (Drinkwater 2005; Mieszkowska et al. 2009). Characterizing the genomic diversity among fish populations, and understanding its relationship to phenotypic variation has become increasingly important in fisheries management and for predicting the response of various ecotypes to environmental fluctuations, such as climatic changes (Neat and Righton 2007; Righton et al. 2010). Earlier studies in Atlantic cod have provided evidence for elevated genomic divergence among populations mainly in respect of four discrete genomic regions, also referred as supergenes, located on linkage groups (LGs) 1, 2, 7 and 12 (Bradbury et al. 2010; Bradbury et al. 2013; Hemmer-Hansen et al. 2013; Karlsen et al. 2013; Berg et al. 2015; Berg et al. 2016; Kirubakaran et al. 2016; Sodeland et al. 2016; Barney et al. 2017; Barth et al. 2017; Berg et al. 2017; Barth et al. 2019; Clucas et al. 2019a; Clucas et al. 2019b; Kess et al. 2019; Puncher et al. 2019). Relationships between these regions and environmental conditions indicates that the region identified on LG01 is associated with strong genetic differentiation between migratory and stationary ecotypes on both sides of the Atlantic Ocean (Hemmer-Hansen et al. 2013; Karlsen et al. 2013; Berg et al. 2016; Kirubakaran et al. 2016; Sinclair-Waters et al. 2017; Kess et al. 2019). This supergene coincides with a double inversion that suppresses homologous recombination in heterozygotes and effectively prevents admixing between co-segregating haplotypes (Kirubakaran et al. 2016). The genomic islands of divergence on LGs 2, 7 and 12 are also found on both sides of the Atlantic Ocean and it has been suggested that they are associated with mean ocean temperatures along the north-south gradient (Bradbury et al. 2010; Bradbury et al. 2013; Berg et al. 2015; Clucas et al. 2019a). Genomic divergence in these regions has also been associated with other environmental factors in studies comparing Baltic and North Sea populations (Berg *et al.* 2015), as well as oceanic and coastal populations in the North Sea (Sodeland et al. 2016). Elevated linkage disequilibrium (LD) detected across the regions on LGs 2, 7, and 12 are likely to have arisen as a result of chromosomal inversions, but high-resolution sequence data showing this and describing the precise locations, sizes and genomic structure underlying these regions has so far been lacking.

Most fish genome sequences have been built from short-read Illumina data, which is a computationally challenging and error prone process especially when the genomes contain extensive repetitive regions. Long-read sequencing technologies provide the means to directly read through repetitive elements and thereby potentially produce much more complete *de novo* assemblies. The recently released gadMor3 assembly (NCBI accession ID: GCF_902167405.1) was developed based on long-read sequence data produced from a NEAC fish and represent a significant improvement over previous gadMor1 and gadMor2 assemblies generated from the same northern population (Star et al. 2011; Torresen et al. 2017). In this paper, we used long-read nanopore data to construct a reference genome assembly for a male Atlantic cod from the southern population of the Celtic Sea and integrated the assembly with linkage data to build high-quality chromosomes sequences. The genome sequence was utilized to detect a potential centromeric repeat sequence differentiating chromosomal morphology and to characterise with high precision the chromosomal rearrangements underlying the notable supergenes on LGs 1, 2, 7 and 12.

## Materials and Methods

### Sample, DNA extraction and sequencing

DNA from a single, male cod (45cm, 1009gm) fished in the Celtic Sea in January (50° 42.16N 07° 53.27W, 110m depth) was extracted from frozen blood using the Nanobind CBB Big DNA kit from Circulomics and sequenced using a PromethION instrument from Oxford Nanopore Technology (ONT). Two sequencing libraries were generated following the ligation protocol (SQK-LSK109, ONT), one using DNA fragments >20kb, size selected using a BUF7510 High pass cassette run on a Blue Pippin (Sage Scientific), and another where no size selection was performed. Both libraries were split in two and each half sequenced successively on the same flow-cell (type R9.4.1) after nuclease flushing according to the Oxford Nanopore protocol (version: NFL_9076_v109_revF_08Oct2018). Combined data yields after quality filtering were 11.2 and 35.5 billion bases for size selected and non-size selected respectively, with median read lengths being 23.3 kb and 4.5 kb. Together this represents approximately 70X long-read genome coverage assuming an Atlantic cod genome size of 670 Mb (as is estimated for gadMor3). Short read data (2 × 250bp) was generated from non-size selected DNA using an Illumina MiSeq instrument. Libraries were prepared using a TruSeq DNA PCR free kit (Illumina) and sequenced in multiple runs to generate 71M read pairs, equalling approximately 35.5Gbp or 50X genome coverage.

### Construction of the gadMor_Celtic assembly

The raw nanopore reads (n=2,868,527) was base-called using Guppy-2.2.3 (https://community.nanoporetech.com) using flip-flop model. Adapters were removed from reads using Porechop v0.2.3, 1 (Wick et al. 2018) and quality-filtered using fastp v.0.19.5.2 (Chen et al. 2018) with mean base quality greater than 7, trimming the 50bp at the 5’ end of the read and removing all reads less than 4000 bp. Multiple initial assemblies applying various parameters were produced using wtdgb2 v2.3 (Ruan and Li 2019). The completeness of all assembled genomes was estimated using BUSCO v3.1.0 (Simao et al. 2015) and applying the actinopterygii (ray-finned fishes) reference gene data set. Two genome assemblies with the relative best values for contig N50, total genome size and BUSCO scores were selected (See File S1) for further quality assessments.

To improve assembly contiguity, contigs showing a sequence overlap of more than 5000bp and similarity >95% were combined using quickmerge (Chakraborty et al. 2016). This consensus assembly was error corrected by performing two successive rounds of processing by Racon v2.3 (Vaser et al. 2017) using only quality filtered nanopore reads. Raw MiSeq reads were quality filtered using Trimmomatic v0.32, before being used by Pilon v1.23 to further improve per-base accuracy in the consensus sequence. Completeness of the final polished contigs was performed as described above using BUSCO.

### Linkage mapping and construction of chromosome sequences

The linkage map was constructed using 9,178 high-quality SNPs (File S2) genotyped in farmed cod (n=2951) sampled from 88 families of the National cod breeding program maintained by Nofima in Tromsø, Norway, and from eight families of the CODBIOBANK at the Institute of Marine Research in Bergen, Norway. The genotypes were produced on a 12K SNP-array created as a part of the Cod SNP Consortium (CSC) in Norway and being used in numerous previous studies (Berg et al. 2015; Sodeland et al. 2016; Barth et al. 2017; Berg et al. 2017; Sinclair-Waters et al. 2017; Knutsen et al. 2018; Kess et al. 2019).The SNPs on this array were carefully chosen to tag as many contigs as possible in the gadMor1 assembly, are thus expected to be well distributed in the genome and builds a good foundation for anchoring sequences to chromosomes. Linkage mapping was performed with the Lep-MAP software in a stepwise procedure (Rastas et al. 2013). First, SNPs were assigned to linkage groups with the ‘SeparateChromosomes’ command using increasing LOD thresholds until the observed number of linkage groups corresponded with the expected haploid chromosome number of 23. Additional SNPs were subsequently added to the groups with the ‘JoinSingles’ command at a more relaxed LOD threshold, and finally SNPs were ordered in each linkage group with the ‘OrderMarkers’ command. Numerous iterations were performed to optimise error and recombination parameters. Following this, sequence flanking each marker was used to precisely position all genetic markers to contigs in the gadMor_Celtic assembly using megablast (Altschul et al. 1990), and thereby associate sequence with linkage groups. This analysis revealed 2 chimeric contigs containing at markers from each of different linkage groups that were selectively ‘broken’ using alignments with the gadMor2 assembly (Torresen et al. 2017). After breakage of the two contigs, linkage information was used to order, orientate and concatenate contigs into 23 chromosomes. Finally, SNPs were positioned in the chromosome sequences using megablast and linkage maps constructed using a fixed order in Lep-MAP to produce the final linkage maps presented in File S2 and File S3.

### Detection of repetitive elements

RepeatModeler version 1.0.8 (Smit et al. 1996) was used to generate a repeat library, subsequently RepeatMasker version 4.0.5 (Smit et al. 1996) was run on the finished gadMor_Celtic with default options to identify the repeats in the genome assembly. For the purposes of detecting putative centromeric sequences, tandem repeats were identified using TandemRepeat finder (TRF) version 4.09 (Benson 1999) with the following parameters: matching weight=2, mismatching penalty=7, indel penalty=7, match probability=80, indel probability=10, minimum score to report=30 and maximum period size to report=500. The output was processed using custom perl and unix scripts to identify repeats specifically containing more than 60% AT, longer than 80 bp, and present in all 23 LGs.

### Gene annotation

Data from various public sources was used to build gene models including (i) 3M transcriptome reads generated using GS-FLX 454 technology and hosted at NCBI’s SRA (https://www.ncbi.nlm.nih.gov/sra/?term=SRP013269), (ii) >250 K ESTs hosted by NCBI (https://www.ncbi.nlm.nih.gov/nucest) (iii) 4.4 M paired-end mRNA MiSeq sequences from whole NEAC larvae at 12 and 35 dph (https://www.ebi.ac.uk/ena, PRJEB25591) and (iv) 362 M Illumina reads from 1 and 7 dph (https://www.ebi.ac.uk/ena, PRJEB25591). To enable model building, MiSeq reads and short illumina reads were mapped to the gadMor_Celtic assembly using STAR v2.3.1z (Dobin et al. 2013), while 454 transcriptome reads were mapped using gmap v2014-07-28 (Wu and Watanabe 2005) with ‘–no-chimeras’ parameter in addition to default parameters. stringtie v1.3.3 (Pertea et al. 2015) was used to assemble the reads into transcript models. Transcript models were merged using stringtie merge (Pertea et al. 2015). Gene models were tested by performing (i) open reading frame (ORF) prediction using TransDecoder (Haas et al. 2013) using both pfamA and pfamB databases for homology searches and a minimum length of 30 amino acids for ORFs without pfam support, and (ii) BLASTP analysis (evalue <1e-10) for all predicted proteins against zebrafish (*Danio rerio*) (v9.75) and three-spined stickleback (*Gasterosteus aculeatus*) (BROADS1.75) annotations from Ensembl. Only gene models with support from at least one of these homology searches were retained. Functional annotation of the predicted transcripts was done using blastx against the SwissProt database. Results from TransDecoder and homology support filtering of putative protein coding loci are shown in File S3.

## Data availability

The datasets generated and used during the current study, gadMor_Celtic, repeat library and all supplementary files files are stored at figshare: doi.org/10.6084/m9.figshare.10252919. The raw nanopore reads used to generate gadMor_Celtic are available at European Nucleotide Archive under accession ID PRJEB35290.

## Results and Discussion

### Genome assembly

The current methodological convention in population genomics is to build genomic tools and interpret results based on the information acquired from one arbitrarily sampled individuals’ reference genome, which is used as a default to represent the whole species. Accordingly, genome assemblies for Atlantic cod have been generated from NEAC, which is a migratory population feeding in the cold waters of Barents Sea. However, with the advent of new, cheaper sequencing platforms and long-read technology it is now possible to develop multiple reference genome sequences representing a broader species diversity. As a contrast to NEAC, we decided here to generate a high-quality reference genome from a male Atlantic cod captured in the Celtic sea, a region representing the southernmost extreme of the Eastern Atlantic distribution (Mieszkowska et al. 2009; Neat et al. 2014) and where cod are likely to be experiencing suboptimal summer temperatures (Neat and Righton 2007). Our gadMor_Celtic assembly was built in a stepwise process involving: (i) the testing of multiple combinations of assembly parameters to generate initial assemblies using wtdgb2 (Ruan and Li 2019); (ii) the merging of contigs from selected initial assemblies into a primary assembly using quickmerge (Chakraborty et al. 2016); (iii) performing multiple rounds of base error correction using Racon (Vaser et al. 2017) and Pilon (Walker et al. 2014); finally (iv) the anchoring and orientation of polished contigs into linkage groups.

The two ‘best’ initial assemblies (see Materials and Methods for details), were similar with regards to their total size (bp), number of contigs, and contig N50 (see Table 1), and their Benchmarking Universal Single-Copy Orthologs (BUSCO) scores of 20-40% indicating a poor content of identifiable reference genes. This last observation likely reflects the fact that they were constructed from nanopore reads alone (which suffer from relatively high rates of substitution and deletion errors; e.g. 13% and 5% respectively (Bowden et al. 2019)) and that the assemblies generated were not corrected with higher quality reads such as those that can be generated from Illumina sequencing (Jain et al. 2018). To improve assembly contiguity, contigs showing a sequence overlap of more than 5kb with >95% similarities were combined using quickmerge. This increased the contig N50 from 6 to 10.4Mb and concurrently reduced the number of contigs. Thereafter, two rounds of error correction were performed. First round used Racon to generate consensus sequences using the 70X nanopore data alone and resulted in a BUSCO score of 66.5%. Second round used Pilon and 50X coverage high-quality Illumina data (16.5Mb paired-end 250 bp reads) and saw the BUSCO genome completeness score increase to 94.2% which is comparable to other high quality fish genomes (e.g. (Chen et al. 2019; Kadobianskyi et al. 2019). The resulting gadMor_Celtic assembly is composed of 1,253 contigs (contig N50=10.5 Mb, average contig length 0.55 Mb) and includes 686 Mb of sequence.

**Table 1.** Assembly statistics. Metrics describing genome statistics of the initial assemblies, the quickmerge assembly, and the final gadMor_Celtic assembly after polishing with nanopore (Racon) and Illumina (Pilon) data.

|  | **Total size**<br>**(bp)** | **Total number**<br>**of contigs** | **Contig N50**<br>**(bp)** | **BUSCO score**<br>**(%)** |
| --- | --- | --- | --- | --- |
| Wtdgb2 assembly 1 | 668,357,526 | 1600 | 6,012,173 | 23.2 |
| Wtdgb2 assembly 2 | 670,278,278 | 1666 | 6,004,590 | 42.4 |
| Quickmerge contigs | 677,547,349 | 1253 | 10,448,158 | Not done |
| Racon polishing | 683,672,734 | 1253 | 10,518,163 | 66.5 |
| **Pilon polishing** | **685,982,295** | **1253** | **10,559,872** | **94.2** |

High-quality linkage maps of densely spaced markers provide the means to reliably anchor genomic fragments (contigs and scaffolds) to chromosomes. If constructed in a large pedigree, and with an adequate number of markers, it may also serve as the backbone for ordering, orienting and concatenating the fragments into chromosome sequences. However, the ability to order and orientate fragments is constrained by the frequency and location of recombination events and thus is limited by the resolution of the map. In this study we used a genetic map consisting of 9,178 SNPs (File S2), constructed in a large pedigree of 2,951 individuals to order and orientate 149 contigs (totalling 643.4 Mb; 93% of assembly) into 23 chromosome sequences. The average number of SNPs per contig was 56.1, with only 12 contigs containing fewer than five SNPs. The high contiguity of the gadMor_Celtic assembly is evidenced by the fact that for one linkage group (LG14), the entire genetic map was correctly captured by a single contig of more than 30 Mb. The total length of the female linkage map (1,662.7 cM) was approximately 1.3 times larger than the male map (1,262.3 cM). The linkage maps were constructed using genotypes from pedigreed samples belonging to families where the large inversions on LGs 1, 2, 7 and 12 were segregating, this led to pronounced gaps in the linkage maps at the boarders of these inversions (see File S4).

### Chromosomal inversions

The detection of extended blocks of LD between SNPs has been used in several studies to define the regions of genetic differentiation on Atlantic cod LGs 1 2, 7 and 12 (Bradbury et al. 2010; Berg et al. 2015; Sodeland et al. 2016; Barney et al. 2017). Large chromosomal inversions have been hypothesized for all four regions but only documented for LG01 (Kirubakaran *et al.* 2016). While regions of extended LD are symptomatic of large polymorphic inversions, no studies have directly compared reference genomes from different cod ecotypes to define and confirm the underlying mechanism, or to locate the genomic regions containing the inversion breakpoints or to define the exact complement of genes they contain. We aligned the recently released gadMor3 assembly (NCBI accession ID: GCF_902167405.1) constructed from a NEAC individual to our gadMor_Celtic assembly using LASTZ (Harris 2007). The gadMor3 assembly was generated following a comprehensive sequencing effort combining long-read sequence data from Pacific BioSciences with various datasets for scaffolding and polishing, and resulted in 1,442 contigs (contig N50=1.015 Mb). Despite being an order of magnitude smaller than our gadMor_Celtic contigs, the gadMor3 scaffolds nevertheless mapped with a high confidence to the assembly and showed that the two assemblies display alternative configurations of inversions for the supergenes on LGs 1, 2, 7 and 12. In most cases, the inversion breakpoints could be described at high resolution because they locate within single nanopore contigs. Exceptions to this were the third breakpoint of LG01 and second breakpoint on LG07 which falls between two gadMor_Celtic contigs (Figure 1).

**Figure 1.**
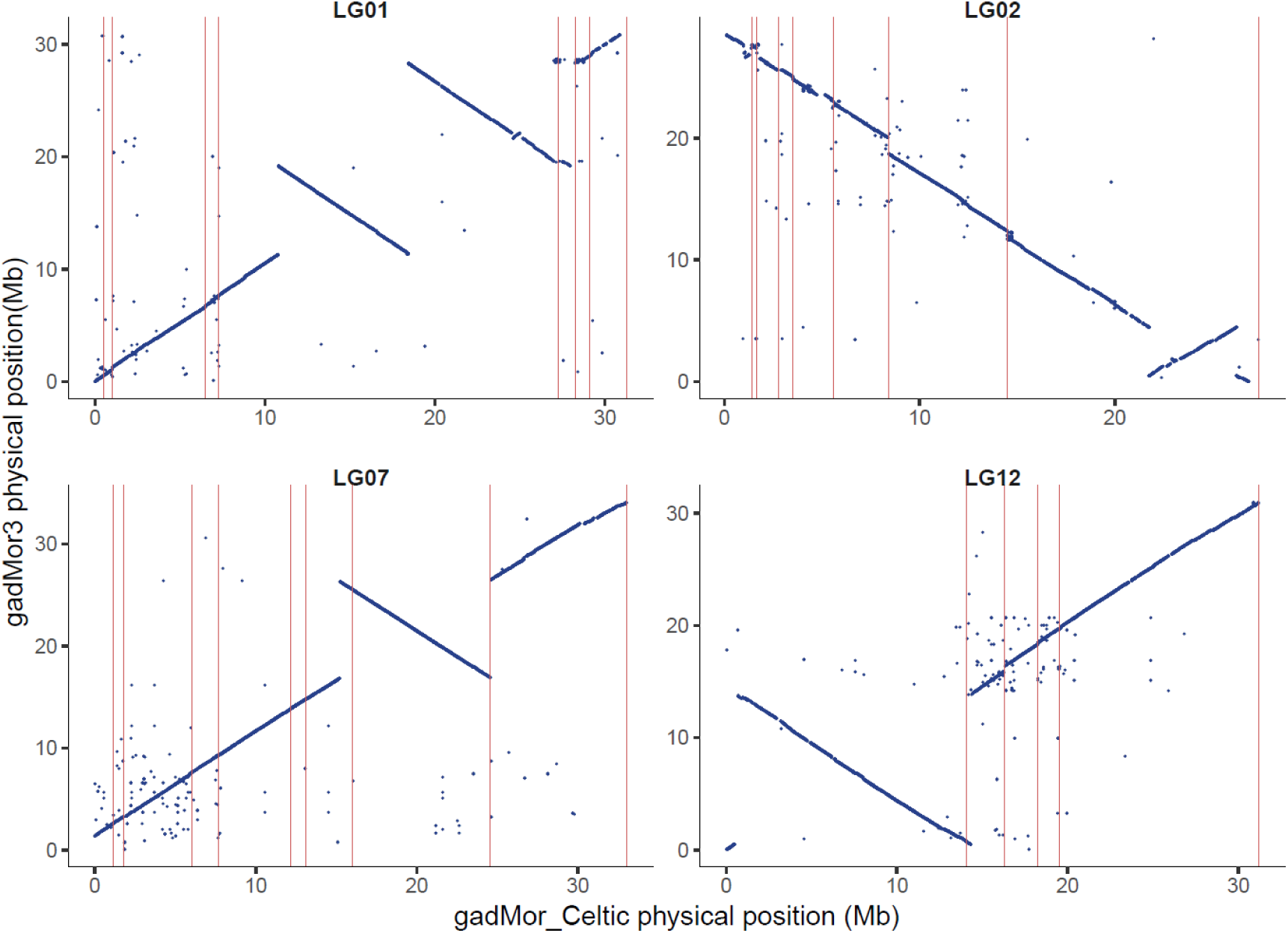
Alignment of gadMor_Celtic (x-axis) and gadMor3 (y-axis) chromosome sequences for linkage groups 1, 2, 7 and 12. Vertical lines (pink) demarcate boundaries of gadMor_Celtic contigs.

A perfect characterization of inversion breakpoints at the sequence level using the gadMor3 and gadMor_Celtic assemblies would require that contigs from both assemblies span the breakpoints and that sequences at the breakpoints would align perfectly with high confidence. As contig structure is not available in the gadMor3 assembly, and genome alignments to some extent were confounded by repetitive sequences, we believe it is appropriate to present the inversion breakpoints as regions, or putative intervals (see Table 2).

**Table 2.** Genomic regions likely containing the inversion breakpoints. A pairwise comparison between gadMor_Celtic and gadMor3 reveals the interval (described as a start and stop coordinates relative to the gadMor_Celtic assembly) for each inversion breakpoint LGs 1, 2, 7, and 12.

| Linkage Group | Putative interval containing breakpoint |  | Size (bp) | Inversion size (Mb) |
| --- | --- | --- | --- | --- |
|  | Start | End |  |  |
| LG01 | 10,782,691 | 10,787,755 | 5,064 | 17.45 |
|  | 18,422,802 | 18,425,099 | 2,297 |  |
|  | 28,225,372 | 28,228,130 | 2,758 |  |
| LG02 | 21,733,338 | 21,733,998 | 660 | 4.51 |
|  | 26,233,253 | 26,238,098 | 4,840 |  |
| LG07 | 15,208,043 | 15,210,043 | 2,000 | 9.37 |
|  | 24,574,346 | 24,575,510 | 1,164 |  |
| LG12 | 493,527 | 635,659 | 142,132 | 13.88 |
|  | 14,330,965 | 14,376,973 | 46,008 |  |

In the gadMor_Celtic assembly the double inversion on LG01 spans a total interval of 17.45 Mb which is slightly larger than our previous estimate of 17.37 Mb (Kirubakaran et al., 2016). Our ability to detect inversions when comparing gadMor3 to the NEAC reference suggests that Celtic cod possess the stationary (as opposed to migratory) ecotype chromosome configuration. An earlier survey of Celtic cod (Neat et al. 2014) showed that while a portion of the population migrate horizontally (from the Celtic sea to the Western English channel) they do not undertake the scale of vertical migration that have been reported for NEAC fish, which have been found at depths of up to 500m (Godo and Michalsen 2000). Instead Celtic cod are typically located at depths of about 100 meters, which is similar to the depth distribution of the stationary populations found around the Norwegian coast (Hobson et al. 2007).

The inversions on LGs 2, 7 and 12 span 4.51, 9.37 and 13.88 Mb, respectively. These sizes are in relatively close agreement to earlier estimations of 5.0, 9.5, and 13 Mb, which were calculated from LD analyses and detection of regions of elevated divergence between populations (Sodeland et al. 2016). In their analysis, Sodeland *et al*. (2016) used the highly fragmented gadMor1 assembly (Star et al. 2011) and a relatively sparse set of 9,187 SNPs to define the regions, both factors that may explain the physical difference between estimates. A more recent study investigated cod populations from the Northwest Atlantic and measured LD amongst almost 3.4M SNPs detected from resequencing data, the LGs 2, 7 and 12 inversions were estimated to be 5.6, 9.3, and 11.6 Mb respectively (Barney et al. 2017). While not identical, these regions and sizes detected in fish from both sides of the Atlantic are remarkably consistent, supporting the hypothesis that these cod have a common ancestral origin (Berg et al. 2017; Sinclair-Waters et al. 2017).

Our analyses suggest the presence of putative inversions in gadMor_Celtic on LGs 6, 11 and 21 (see Figure 2) which, to the best of our knowledge, have not been reported elsewhere. The inversions are smaller (1.4, 0.6, 1.78 Mb, respectively) than the rearrangements comprising the supergenes on LGs 1, 2, 7 and 12.

**Figure 2.**
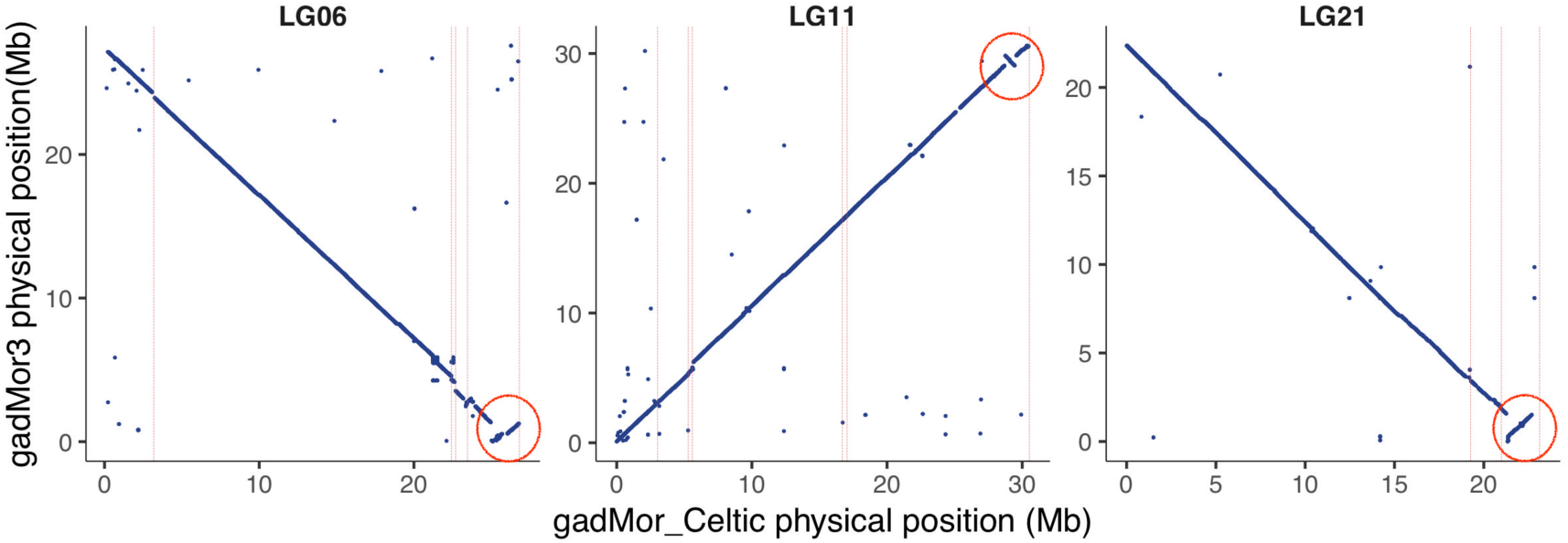
Putative inversions detected on LGs 6, 11 and 21.

### Annotation of gene content and repetitive elements

There is a growing body of evidence that chromosomal inversions in fishes can capture multiple adaptive alleles and therefore act as supergenes (for example (Jones et al. 2012; Pearse et al. 2018; Pettersson et al. 2019). Defining the gene content and identifying genetic variation within these chromosomal inversions is an important means for investigating how changes in genome organization may lead to phenotypic and adaptive divergence. Utilizing available transcript data we predict 14,292 genome wide gene models with 735, 236, 343 and 452 gene models predicted in inversions on LGs 1, 2, 7 and 12 respectively (File S5). In the context of a north versus south contrast (i.e. NEAC vs Celtic) the polymorphic haemoglobin *Hbβ1* gene deserves special mention since there is good evidence for temperature-associated adaptation (Frydenberg et al. 1965; Andersen 2012). Although the haemoglobin gene maps to LG02 it is, however, located outside the inversion (approximately 3 Mb upstream) which raises questions about the mechanism maintaining its association with temperature. To document repeats in gadMor_Celtic we created a repeat library using RepeatModeler (Smit et al. 1996) which, when used with RepeatMasker (Smit et al. 1996) saw almost one third of the genome (32.26%) classified as repetitive.

### Potential centromere structure and organization

Centromeres contribute to the physical linking of sister chromatids during meiosis and their location within a dyad is important for defining the chromosomal morphology (or chromosome classification) used in karyotyping studies (e.g. metacentic, acrocentric, etc). Centromeres can be relatively large and usually contain a lot of repetitive, but poorly conserved sequences (Melters et al. 2013). Searching for known centromere repeats (Melters et al. 2013) in gadMor_Celtic assembly failed to reveal any convincing hits. We therefore used TandemRepeat finder (TRF) (Benson 1999) to scan the assembly for seqences meeting characteristics typical of centromeric repeats; specifically containing more than 60% AT, longer than 80 bp, and present in all 23 LGs. We detected a 258bp sequence composed of two identical and similarly oriented 88bp repeats (one at each end) separated by an 82bp interveining sequence (see File S6 for details). This expected centromeric repeat appeared 806 times (with more than 95% identity) across the genome and was found on all LGs. The location of this repeat was compared to the genetic map profiles for all 23 linkage groups (File S4). We reasoned that regions of reduced recombination likely contain, or are close to, the centromere and should therefore coincide with the mapping of the centromeric repeat sequence. For most linkage groups, there was a convincing overlap between these two metrics. Most evidently, all four LGs (2, 4, 10 and 12) showing clear sigmoidal linkage profiles characteristic of a metacentric chromosome (Ghigliotti et al. 2012), contained expansion of the centromeric repeat sequence within the region of repressed recombination in the middle of the linkage group (see Figure 3 for example).

**Figure 3.**
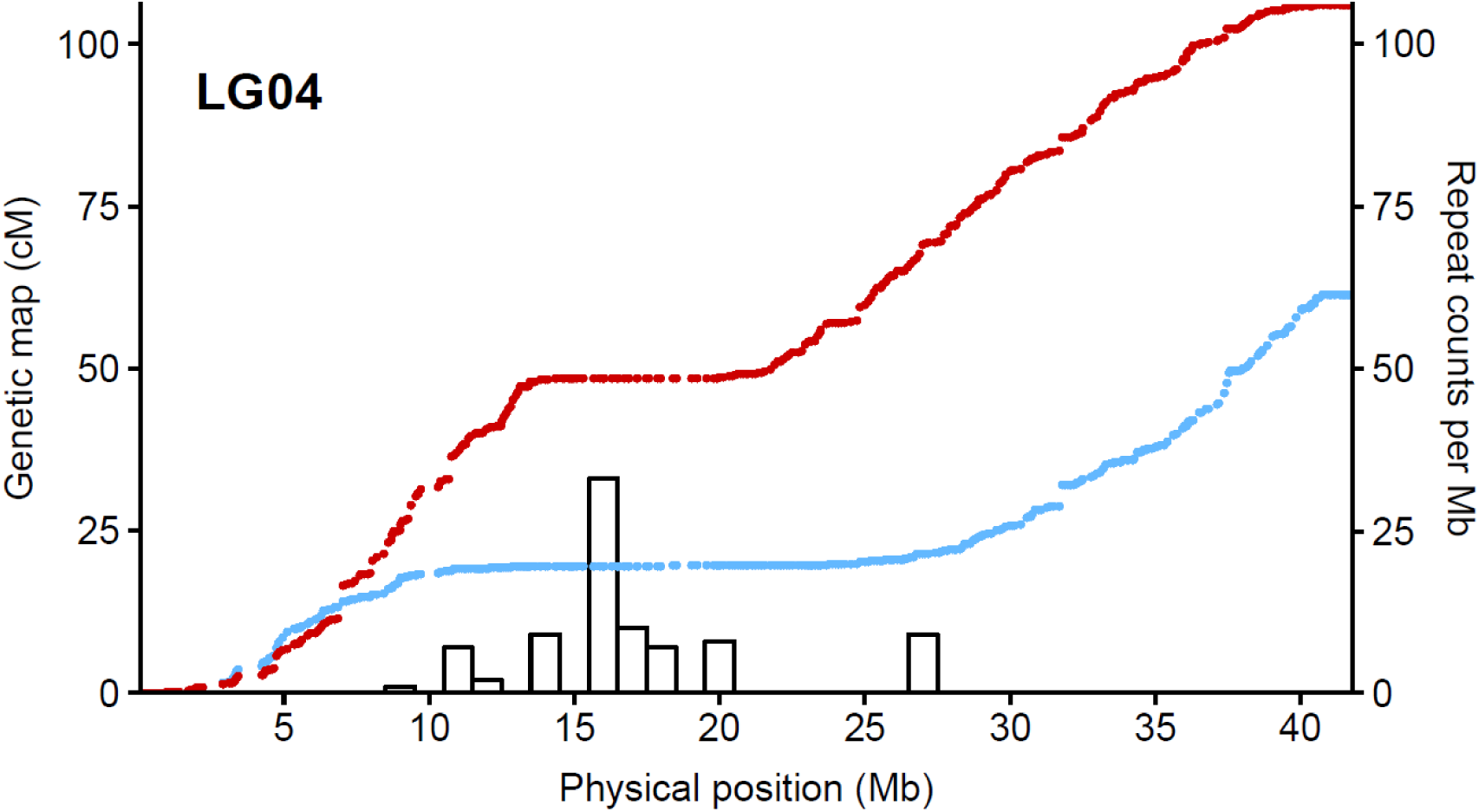
Position of potential centromere related sequence on LG04. Collinearity between LG04 genetic maps for males (red) and female (blue) and the frequency of a 258bp tandem repeat structure (histogram) predicted to be related to centromeres.

In this paper we used nanopore sequencing to generate a chromosome-level genome assembly from a male Atlantic cod captured in the Celtic Sea. Cod from this region experience high, possibly suboptimal summer temperatures, and consequently this sample represents a contrast to the current genome assemblies generated from NEAC population sampled from the considerably colder Barents Sea. By generating this new assembly, and comparing it against the gadMor3 assembly, we were able to characterize the population specific chromosomal rearrangements associated with four notable supergenes displaying pronounced divergence between them. Pairwise comparison of the two genomes also revealed additional putative rearrangements on LGs 6, 11 and 21, which has not been reported before. Identification and mapping of the centromeric repeat enabled by the new high resolution gadMor_Celtic assembly, combined with linkage maps, were used to study chromosomal morphology and reliably identify four characteristic metacentric chromosomes in Atlantic cod.

## Supplementary Material

File S1: wtdbg2 parameters used to generate the two initial genome assemblies.

File S2: Linkage map of gadMor_Celtic: SNPs, position in gadMor_Celtic, genetic linkage of male, female in centimorgan (cM) and SNP flank sequence from gadMor1 (NEAC).

File S3: Predicted function of open reading frames were found with TransDecoder and homology search using blastp against zebrafish and stickleback protein databases.

File S4: Plots showing collinearity between genetic maps for males (red) and female (blue) and the frequency of a 258bp tandem repeat structure (histogram) predicted to be related to centromeres in all 23 chromosomes.

File S5: This contains the list of genes and its positions in LGs 1, 2, 7 and 12.

File S6: The putative 258bp centromere repeat sequence.

## Acknowledgements

The authors are grateful to Mr Brendan O’Hea at the Fisheries Ecosystems Advisory Service for providing the fish samples, and to our colleagues in the Cod SNP Consortium (CSC; a collaboration between CIGENE, CEES, IMR and Nofima) from where the genotypes used for the linkage analyses were derived. Funding for T.G.K. was provided by the Norwegian University of Life Sciences (NMBU).

